# Maximizing Ecosystem Services Provided to the New Oil Crop *Brassica carinata* Through Landscape and Arthropod Diversity

**DOI:** 10.1101/724203

**Authors:** Shane Stiles, Jon Lundgren, Charles Fenster, Henning Nottebrock

## Abstract

Prairies, once spanning the Upper Midwest, have now largely been replaced by agriculture. The lack of resources available to pollinators in agricultural fields and practices commonly employed has led to a decline in insect diversity. To enhance sustainable practices, we must better understand how ecosystem services such as pest control and pollination services provided by a diverse insect and pollinator community scale to current farming practices as related to crop yield and how landscape features may positively contribute to insect and pollinator diversity. We examined how landscape heterogeneity relates to insect and pollinator diversity, as well as how insect and pollinator diversity relates to crop yield across common farming practices. We planted 35 single acre sites of *Brassica carinata,* a generalist flower possibly capable of supporting a diverse insect community. We randomly assigned each site with a combination of three common farming practices: tilling (yes/no), added honey bee hives (yes/no), and treatment with systemic neonicotinoids (yes/no). Insect and pollinator diversity and the surrounding landscape at multiple spatial scales were calculated. We observed a significant positive relationship between insect (and pollinator) diversity with yield in the absence of any farming practice. All farming practices will increase yield. However, farming practices alter the relationship between yield and diversity. The addition of seed treatment or tillage negates the relationship between insect (and pollinator) diversity with yield. Seed treatment alone results in a flat relationship between diversity and yield for all insects and a negative relationship for pollinators. Increased landscape heterogeneity results in a positive relationship between insect diversity at the 1000 m scale and pollinator diversity at the 3000 m scale, suggesting large-scale heterogeneity contributes to overall insect diversity. Our results show that increasing large-scale landscape heterogeneity increases diversity serving as a substitute for common farming practices such as application of pesticides, tilling, or bee hives. Increased heterogeneity could save farmers from the input cost of treatment or tillage, by way of increased insect diversity, while still providing similar yields.

## Introduction

Intensive agriculture has replaced prairies as the primary land use within the Upper Midwest (McGregor, 1986). These lands, once rich in biodiversity, have been converted to landscapes dominated by corn and soybean, providing minimal nectar and pollen forage to arthropods such as pollinators (Smart et al. 2016). Additionally, practices employed by farmers to maximize yield such as tilling (McLaughlin and Mineau 1995), pesticide treatment (Rundlöf et al. 2015), and added honey bee hives (Mallinger et al. 2017), has led to a decline in insect and pollinator diversity (Kearns et al. 1998). Landscape heterogeneity or the diversity of land uses at the landscape scale has declined with agricultural intensification (Benton et al. 2003). These changes have altered ecosystem services provided by arthropods to humans, including pollination services, natural pest control and biodiversity. However, appropriate management of ecosystem services may ameliorate many of the negative impacts of agriculture (Power 2010). An important gap in our understanding of pollination services and natural pest management is that we do not know how these ecosystem services scale to current farming practices as they relate to crop yield. Biodiversity plays an essential role in agricultural landscapes, which requires quantifications of how biodiversity impacts the productivity (Scherr and McNeely 2008). We are lacking information on the functional role of biodiversity for yield productivity and how this relationship is impacted by current farming practices.

Ecosystem services such as pollination services play an essential role in agricultural landscapes. In particular, wild pollinators can contribute more to crop yield than domesticated honey bees (Garibaldi et al. 2013, Mallinger and Gratton 2015, Lindström et al. 2016). However, land use change has had an impact on pollination services provided to crops by decreasing pollinator diversity (Grab et al. 2019). Additionally, there are instances of agriculture favoring agrobiont species that compensate for the services provided by pollinators displaced with intensification (Mogren et al. 2016). Agricultural intensification generally jeopardizes wild bee communities and hence the pollination services they offer to crops (Klein et al. 2007). Stabilizing biodiversity by increasing landscape heterogeneity could enhance pollination services and natural pest control by reducing the risk to instability (Jackson et al. 2005).

Farming practices occur within a broader agricultural ecosystem that in turn influences biodiversity such as insect diversity (Fahrig et al. 2011). For example, and our focus here, increased landscape heterogeneity promotes the maintenance of biodiversity, and consequently ecosystem services (Loreau et al. 2003, Tscharntke et al. 2005). As landscape heterogeneity increases, the abundance and diversity of natural enemies increases, stabilizing pest communities while increasing insect diversity (Bianchi et al. 2006, Chaplin-Kramer et al. 2011, Veres et al. 2013). Natural areas adjacent to nectar-providing crops can also increase the abundance of wild bee species (Holzschuh et al. 2013). Increasing insect diversity could both increase pollination services and mediate pest populations by heterogeneous landscapes with sufficient supply of floral resources. Thus, to have a better understanding of how insect and pollinator diversity is related to crop yield we need to consider the landscape context in which farming inputs occur. Linking landscape heterogeneity and farming practices with insect/pollinator diversity and yield is an area in which more attention is required.

Common farming practices such as treatment with neonicotinoids, tillage, and added honey bee hives (henceforth referred to as common farming practices) could increase crop yields. Tillage can have a positive overall effect on crop yield under some circumstances, but its benefits are crop and context dependent, for example it has a mild to negative effect on *Brassica napus* yields (Arvidsson et al. 2014). Additionally, added honey bees do not always fully replace the yield benefits offered by wild pollinators (Garibaldi et al. 2013) yet are used in 90% of all commercial pollination (Genersch 2010). Neonicotinoids have been effective in decreasing pests of differing crops (Elbert et al. 2008, Perkins et al. 2018, Lahiri et al. 2019). However, we have little understanding of how these common farming practices interact with insect diversity in terms of crop yield. The possibility of increased yields through a combination of anthropogenic inputs and ecosystem services would be of great interest to producers.

We use the oilseed mustard *Brassica carinata* (carinata, Brassicaceae), a mass flowering crop, to examine the relationship between insect/pollinator diversity and yield within a broader agroecosystem landscape context. This crop was chosen because it is currently being developed as a biofuel, for aid in phytoremediation, and as a feedstock (Cardone et al. 2003, Anjum et al. 2012, Licata et al. 2018). It is increasingly being used as a winter annual cover crop in the Southeast USA and may be an important component of the weed/ fallow rotation in the Upper Midwest. In addition, production of carinata occurs throughout the world including South America and Australia. Carinata has a flower indicative of generalized pollination, confirmed by our own observations (Nottebrock et al. unpublished) visited by many pollinating species. Carinata also shares a similar pollination system with other flowering crops in the region, especially canola. There is also a need to study ecological properties of this crop as it becomes more common. Cultivation of this crop could have ecological benefits as a provider of resources to pollinators, similar to what is found with its relative *Brassica napus* (e.g., canola) or it could dilute pollinators at the landscape scale (Westphal et al. 2003, Holzschuh et al. 2013, Thom et al. 2016). More research is needed on the effects of mass flowering crops to pollinator health.

Our study explicitly quantifies the effect of insect and pollinator diversity on yield across a range of common farming practices as well as landscape determinants of insect and pollinator diversity by addressing the following questions. First, is there a relationship between insect/pollinator diversity with carinata yield? Second, are insect/pollinator diversity yield effects modified by the common farming practices of tilling, seed treatment with neonicotinoids, and the addition of honey bee hives? Third, is there a relationship between landscape and insect/pollinator diversity found within our plots, and if so, at what scale? Fourth, can the range of common farming practices mentioned above influence the insect/pollinator relationship with the landscape? Our overall goal is to evaluate farming practices and landscape heterogeneity to determine the strongest predictors of carinata yield and insect/pollinator diversity.

Our questions become especially relevant in the context of global food security. Climate change is expected to impact the access, availability, and prices of crops in the future (Schmidhuber and Tubiello 2007) due to the shifts in insect community habitat and the northward expansion of planting zones. Knowledge of these effects could be used for future prediction of yield under an altered climate. Furthermore, despite the tension between crop and grazing practices at the traditional juncture of long- and short-grass prairies (Collins et al. 1998), the northern Great Plains has been understudied relative to other regions in terms of our understanding of the nexus between invertebrate diversity, landscape heterogeneity and crop yield (Dainese et al. 2019).

## Materials and Methods

### STUDY DESIGN

In 2017 and 2018 we planted carinata in 19 × 1-acre sites and in 16 × 1-acre sites, in Brookings and Kingsbury Counties, South Dakota, respectively (appendix S1). To measure how common farming practices might interact with insect diversity and carinata yield, we employed a 2 × 2 × 2 factorial design. Carinata of unknown variety, acquired from Green Cover Seed in Bladen, NE, was planted in May-June of 2017 and May of 2018 with a grain seed drill. Seeding rate was 9 kg/ha (or 8 lbs./acre) and row spacing was 19 cm (7.5 in) apart. We chose sites surrounded by varying degrees of heterogeneity, ranging from many different land uses with an irregular distribution to few land uses with a more even distribution using the classification of land use types from cropscape (USDA). All sites were randomly assigned a combination of three treatments: seeds treated with Poncho 600^®^, a systemic neonicotinoid containing 48% clothianidin (yes, no), four added honey bee hives (yes, no), and tilled (yes, no). Overall, 17 sites were treated with clothianidin, 18 sites had honey bee hives, and 22 sites were tilled. Honey bee hives were deployed soon after planting directly adjacent to the carinata fields. At deployment, hives had approximately 8 frames of bees, received no sugar supplemental feeds, and were actively managed over the season to facilitate hive growth. A table of all treatments received by each site can be found in appendix S2. All sites were treated with Roundup^®^ prior to carinata germination and all sites were treated with a grass herbicide about one month post emergence, Medall II^®^ in 2017, and Poast^®^ in 2018.

### INSECT COLLECTIONS

Sweep samples were used to quantify insect diversity. A 30 m transect was randomly established in each field on each sample date. We averaged 3 and 4 samples per site in 2017 and 2018, respectively. The net was 15 inches in diameter. Samples were collected parallel to the transect. Once insects were collected they were frozen until identification. Across both growing seasons, a total of 142 sweep samples were collected and identified. Insects were observed under an Olympus^®^ microscope made by Diagnostic Instruments Inc and using GSQH10X/22 oculars. Insects were identified to family, then to morphospecies within family for both years using Borror and Delong’s Study of Insects 7^th^ edition, as well as bugguide.com. Pollinators were assigned according to a literature search of each family.

### HARVEST

Carinata grows to 280 cm tall and produces up to 200 flowers per season. Every silqueous fruit contains 10-20 seeds with a high protein and oil content (Basili and Rossi 2018) and thus has the appearance of a robust canola plant. We harvested plants when the fruits were brown and almost dehiscent. To determine plant yield, five randomly selected carinata plants per 1 m^2^ were cut at ground level and placed into individual paper bags. The remaining carinata plants in the quadrat were harvested and placed into a larger bag. All plants were dried in a drying oven until a consistent weight was reached. To perform yield estimation for each site, every random plant was weighed, the total fruits per random plant were counted, and five fruits from each random plant were selected. The seeds of each fruit were weighed and the number of viable and aborted seeds was counted to estimate yield per plant. Average weights and yields of each focal plant within a quadrat were used to estimate the yield of that quadrat based on its weight. To estimate site yield, we used a linear regression model that predicts the number of seeds with plant weight described below.

### ANALYSES

#### Yield Calculation

For all statistical analyses we used R version 3.5.1 (R Core Team, 2018). Calculations of site yield included seed number and weight information of all harvested focal plant individuals per 1 m^2^ and site. Every site was used as an experimental unit. We performed a linear mixed effect model with R package lme4 (Bates et al. 2015) using number of fruits as the response variable and plant weight as the predictor variable. In addition, the average weight of an individual seed per site and the site itself were used as random effects to correct for differences between the sites and ripeness of the fruits. Model predictions were then used to calculate the seed weight per m^2^ based on the biomass weight and yield of each focal plant for each quadrat. In other words, the yield of each quadrat was determined by the yields of each focal plant. Finally, the site yield was predicted by the mean yield of five quadrats per site and multiplied by 10,000 (the number of square meters in a hectare) to estimate the yield of each site in units of kg/ha. Yield calculations can be found in data S1.

#### Landscape Heterogeneity Quantification

To estimate landscape heterogeneity, we used a Trimble GeoXH 2005 dGPS with up to 10 cm accuracy to record the center of each site as a data point. We then obtained a raster file (matrix of pixels organized into a grid in which each pixel contains a colored value representing a specific land use) of 2017 and 2018 USDA Cropscape data. Vector shapefiles were then created at three radii from each of our site points (500 m, 1000 m, 3000 m). Vector files were created in QGIS version 2.18.9. The proportional land use indices were calculated by clipping the raster surrounding every individual site to its appropriate vector radii diameters, and then using the GRASS ‘r.report’ feature located inside QGIS to determine the number of pixels corresponding to each land use. Shannon diversity (H) of the landscape surrounding each site was calculated using the ‘vegan’ community ecology package (R package version 2.4-6). Landscape Quantification code can be found in data S2.

#### Insect and Pollinator Diversity Calculation

The diversity of insects and pollinators were calculated using the Shannon index (data S3) estimated with the ‘plyr’ package in R. The Shannon index is calculated using the following formula:

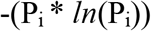

where P_i_ is the sum of the proportions of each species, and *ln* is the natural log. Qualitatively similar results were observed for the relationship of landscape heterogeneity and yield with the separate components of the Shannon index.

#### Relationship between insect/pollinator diversity and carinata yield

To compare yield with insect and pollinator diversity we used linear mixed effect models with R package lme4 (Bates et al. 2015) with year as the random intercept effect. To meet assumptions of a normal distribution for the analyses, yield was natural log +1 transformed. Our three farming practices and diversity metrics are fixed effects as shown by the following formula:

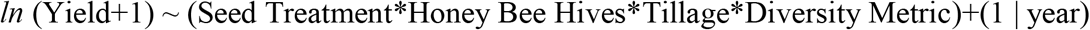

The model includes all main and interactive effects. Our models were then simplified using stepwise-backward variable selection (Crawley 2013). We tested for the inclusion of non-significant main and interaction effects using likelihood-ratio tests with chi-square as a criterion for model assessment. The best overall model included all factors through the three-way interactions even though several of the two-way were not significant. The two-way interactions were included because they contribute to the three-way interactions. All models can be found in data S4. The main and interaction effects are summarized as coefficient effect sizes. Each farming practice effect reflects its increase in yield, holding all other variables constant. Each farming practice effect reflects the absence of all other farming practices due to their categorical nature and holding constant the continuous diversity metrics. Interaction effects are added to the sum of the main effects that comprise the interaction. If an interaction effect is absent we must assume that the main effects are additive. To translate the effect size into increased yield in kg/ha raise *e* to the effect size (*e*^effect^ ^size^).

We did not include landscape heterogeneity into models that predict yield through insect/pollinator diversity for two reasons. First, we expect landscape heterogeneity to affect yield through diversity and so we focus on the relationship between landscape heterogeneity and diversity. Furthermore, in the analyses presented below, we found no relationship between farming practices and insect/pollinator diversity, suggesting that landscape heterogeneity is a major determinant of insect/pollinator diversity in the carinata sites.

#### Relationship between insect/pollinator diversity and landscape heterogeneity

To compare insect/pollinator diversity to landscape heterogeneity we used a model much like the previous model. The three landscape scales were multiplied by the three farming practices and again, simplified using stepwise backward variable selection. Best model assessment was determined as above. The diversity metric was natural log transformed. The formula is shown below:

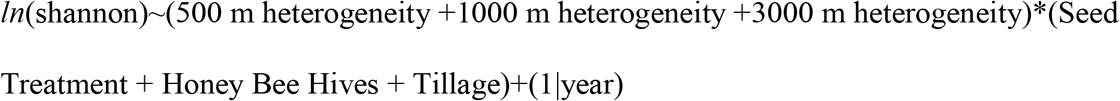

#### Relationship between landscape heterogeneity and yield

To compare the relationship between yield and landscape qualities, yield was again natural log +1 transformed and compared to the three landscape scales and farming practices, using year as a random intercept effect as shown below:

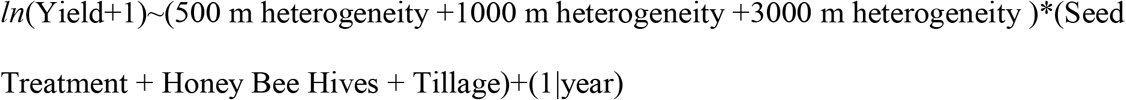

Best model assessment was determined as above.

## Results

We present results for 6 analyses: 1) yield predicted by insect diversity across farming practices, yield predicted by pollinator diversity across farming practices, 3) insect and pollinator diversity across farming practices, 4) insect diversity predicted by landscape heterogeneity across farming practices, 5) pollinator diversity predicted by landscape heterogeneity across farming practices, and 6) yield predicted by landscape heterogeneity across farming practices. We only present model output for the best models confirmed by likelihood ratio tests. Because the analyses are conducted on yield data that have been natural log transformed, the results represented in figures are plotted on a *ln* scale. To make the results more comprehensible, we provide estimated ranges of the yield in kg in the text presented below. The significant effects of main factors and their interactions are presented as effect sizes (± one standard error) of yield (kg/ha). Insect and pollinator diversity are presented separately. For purposes of simplicity, interaction strengths alone are presented without the combined effects of main interactions. Yield is presented in appendix S3: tables S1 and S2.

### Model 1) INSECT DIVERSITY AND YIELD

#### Best model for carinata yield including insect diversity and all farming practices

The minimal adequate model includes two significant three-way interactions with the respective two-way interactions and main effects: the three-way interaction between seed treatment, tillage, and insect diversity is positive with an effect of 2.25±0.98 (adding 54.69 kg/ha) (LRT: χ^2^_1DF_ = 7.6197, p = 0.006). The three-way interaction between seed treatment, added honey bee hives, and insect diversity is also positive adding 64.84 kg/ha with an effect of 2.17±0.93 (LRT: χ^2^_1DF_ = 7.9696, p = 0.005) (Fig. 1, Appendix S3: Table S1). In the following we describe all other significant interaction and main effects.

**FIGURE 1:**
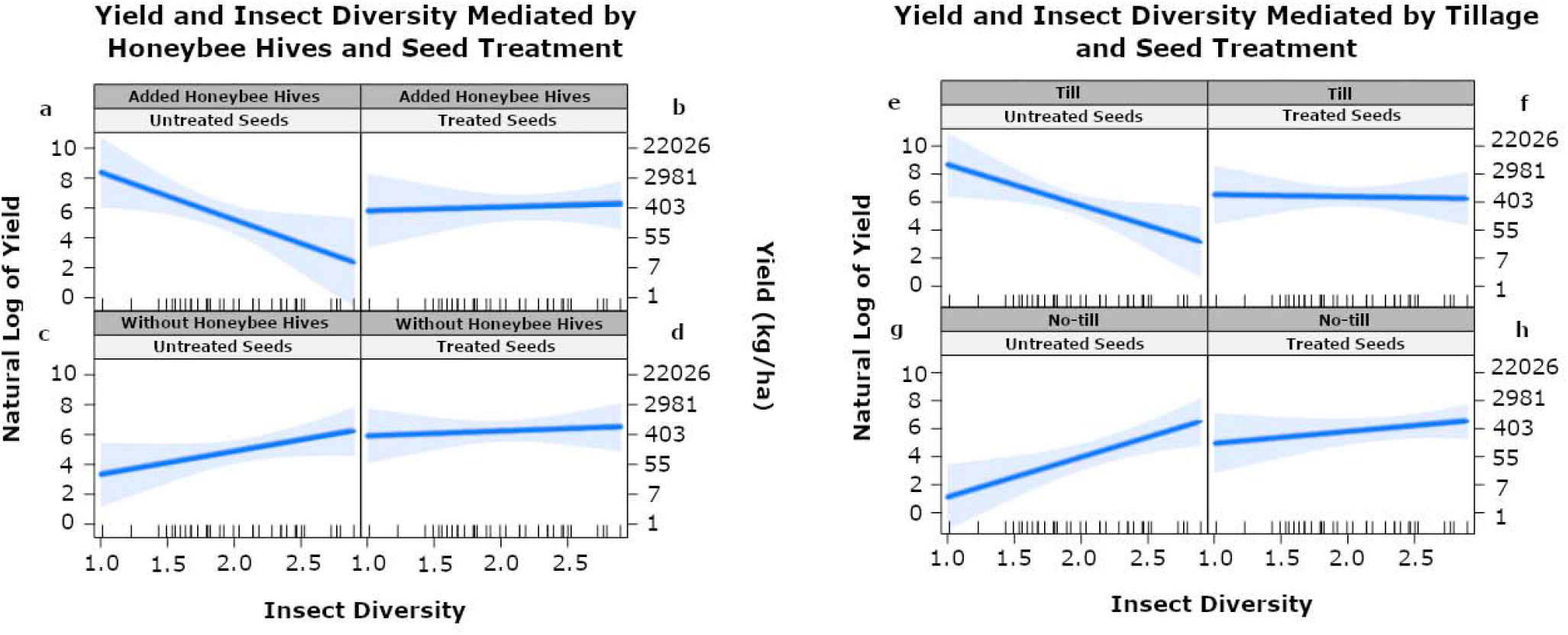
Linear mixed effect analysis of main effects, 2-way, and 3-way interactions between tillage and seed treatment (a-d) and treatment and added honey bee hives (e-h), with insect diversity on *Brassica carinata* yield in eastern South Dakota, 2017-2018. Ticks on the x-axis represent insect diversity of each site.

#### Two-way interactions of farming practices and insect diversity on yield

All two-way interactions of the three farming practices with insect diversity have a significant negative interaction on yield. Seed treatment has the smallest interaction with diversity and an effect size of −1.87±0.72 adding 10.45 kg/ha while tillage has the strongest interaction with diversity at an effect size of −2.71±0.76 adding 15.16 kg/ha. In other words, farming practices are not additive with insect diversity, meaning that increasing insect diversity in the presence of any of our common farming practices will not result in increased yield (Fig. 1, Appendix S3: Table S1).

#### Two-way interactions among farming practices on yield

Added honey bee hives and tillage are the only farming practices that show a significant interaction on carinata yield independent of insect diversity. This interaction has a strong negative relationship with an effect size of −3.63±0.83 or increasing yield by 6.32 kg/ha (LRT: χ^2^_1DF_ = 21.8007, p < 0.001). That is, the combination of added hives and tilling is not additive, but results in a yield roughly the same as either farming practice alone. Neither seed treatment × tillage, nor seed treatment × added honey bee hives had interaction effects that were significant, but they were kept in the simplified model after performing a chi-square test that indicated better model performance with their inclusion.

#### Main effects of farming practices on yield

All main effects, seed treatment, added honey bee hives, tillage, and insect diversity, have a significant positive effect on yield when all other variables are controlled. However, the relationship between these factors and yield change according to farming practices. Tillage has the strongest effect on yield, adding 23.32±1.94 kg carinata seed/ha (p < 0.001), while seed treatment has the weakest at 6.96±2.03 kg/ha (p=0.013). Insect Shannon diversity and honey bee treatment have intermediate effects on yield; there were 9.80±1.06 kg/ha (p = 0.002) for every unit increase in Shannon, and 10.17±2.09 kg/ha when honey bee hives were adjacent to the carinata fields (p = 0.005) (Fig. 1, Fig. 2, Appendix S3: Table S1).

**FIGURE 2:**
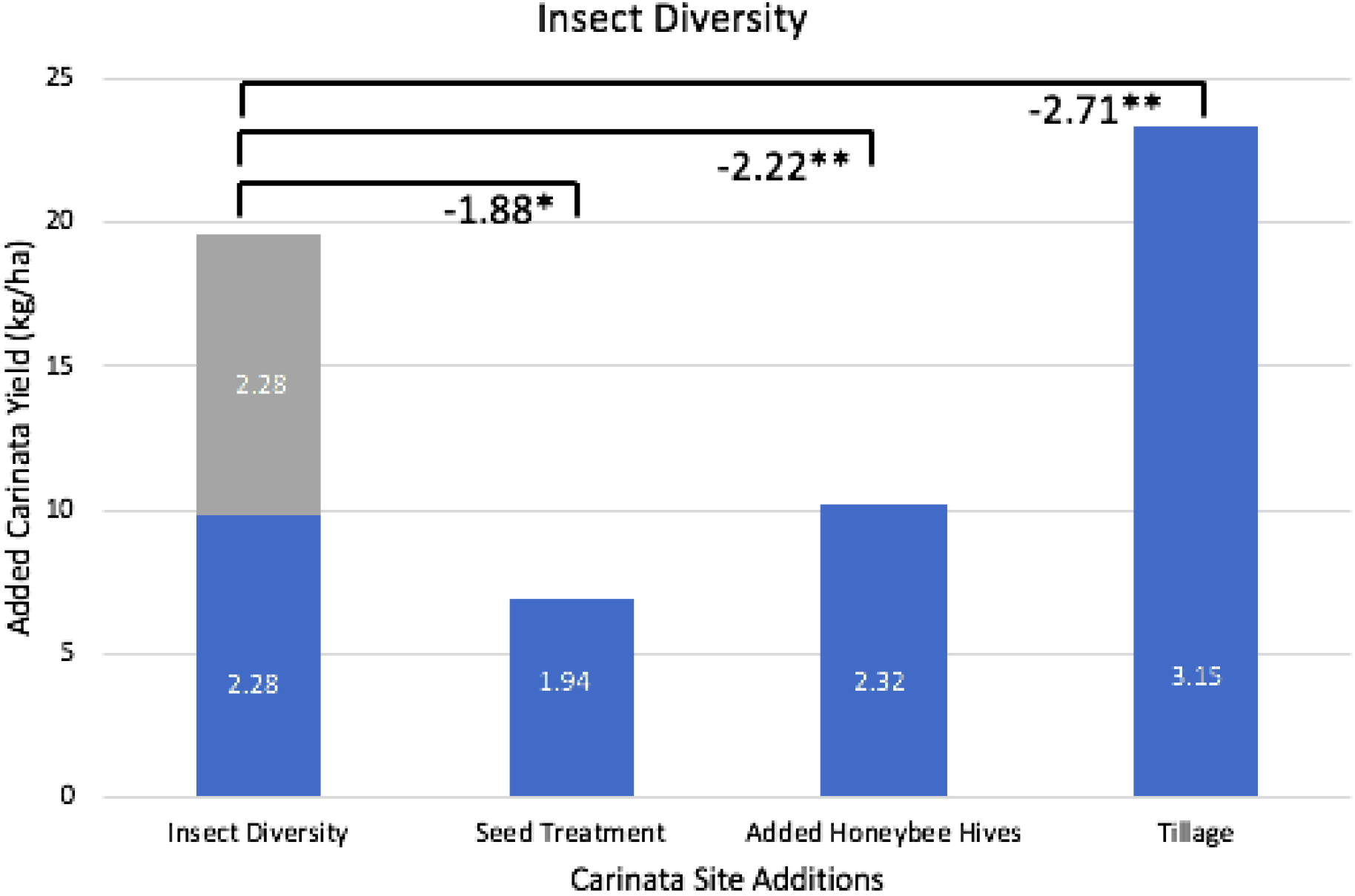
Best model estimates of carinata yield added by each farming practice and Shannon insect diversity shown as main effects. The estimates are based on the fitted values from the model including interactions of farming practices and insect diversity. The reduction or increase of yield depending on interactions with common farming practices is plotted in Fig 1. The blue Insect Diversity bar represents an increase from 1-2 on the Shannon index, while the gray bar represents a jump from 2-3. Numbers inside the bars represent effect sizes before back-transformation. Brackets indicate the negative interactions between insect diversity and the common farming practices. *P* < 0.05; * *P* < 0.05; ** *P* < 0.01; *** *P* < 0.001.

### Model 2) POLLINATOR DIVERSITY AND YIELD

#### Best model for carinata yield including pollinator diversity and all farming practices

The minimal adequate model includes two significant three-way interactions with the respective two-way interactions and main effects: Seed treatment, tillage, and pollinator diversity interact with an effect of 2.91±0.78 (6 kg/ha) (LRT: χ^2^_1DF_ = 16.367, p < 0.001). Added honey bee hives, tillage, and pollinator diversity has an effect of 2.56±0.69 (4.20 kg/ha) (LRT: χ^2^_1DF_ = 15.902, p < 0.001) (Fig. 3, Appendix S3: Table S2).

**FIGURE 3:**
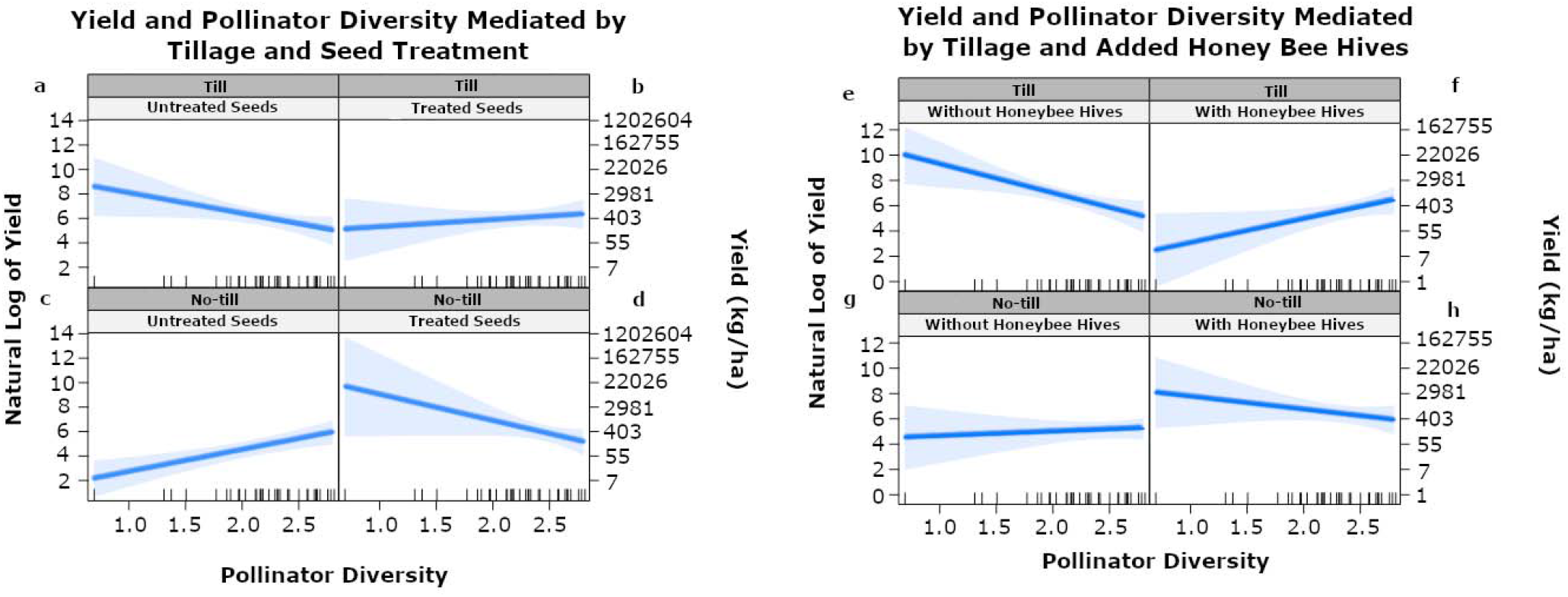
Linear mixed effect analysis of main effects, 2-way, and 3-way interactions between tillage and seed treatment (a-d) and tillage and added honey bee hives (e-h), with pollinator diversity on *Brassica carinata* yield in eastern South Dakota, 2017-2018. Ticks on the x-axis represent pollinator diversity of each site.

#### Two-way interactions of farming practices and pollinator diversity on yield

Seed treatment and pollinator diversity interact with an effect of −1.83±0.56 (2.48 kg/ha). Tillage and pollinator diversity interact with an effect of −2.72±0.54 (2.16 kg/ha). Again, these values indicate that the effect size is lower than the additive values of their main effects (Fig. 3, Appendix S3: Table S2).

#### Two-way interactions among farming practices on yield

All two-way interactions between farming practices and pollinator diversity on yield are negative, suggesting that farming practices interfere with the relationship between pollinator diversity and yield. The interaction between seed treatment and added honey bee hives is not significant, but is withheld in the final model in accordance with the chi-square test. Seed treatment and tillage significantly interact with effect sizes of −1.69±0.62 (10.35 kg/ha). Added honey bee hives and tillage also interact with an effect of −2.78±0.62 (3.06 kg/ha).

#### Main effects of farming practices and diversity on yield

Seed treatment, added honey bee hives, tillage, and pollinator diversity all have a significant positive effect on yield controlling for all other variables. Tillage has the strongest effect on yield with an effect of 2.39±0.48 (10.91±1.61 kg/ha) (p < 0.001), while pollinator diversity has the weakest at 1.10±0.21(3±1.2 kg/ha) (p < 0.001). Seed treatment and added honey bee hive effects are of intermediate strength effect sizes of 1.64±0.5 (5.15±1.64 kg/ha) (p = 0.003) and 1.51±0.42 (4.52±1.52 kg/ha) (p = 0.002) respectively (Fig. 3, Fig. 4, Appendix S3: Table S2).

**FIGURE 4:**
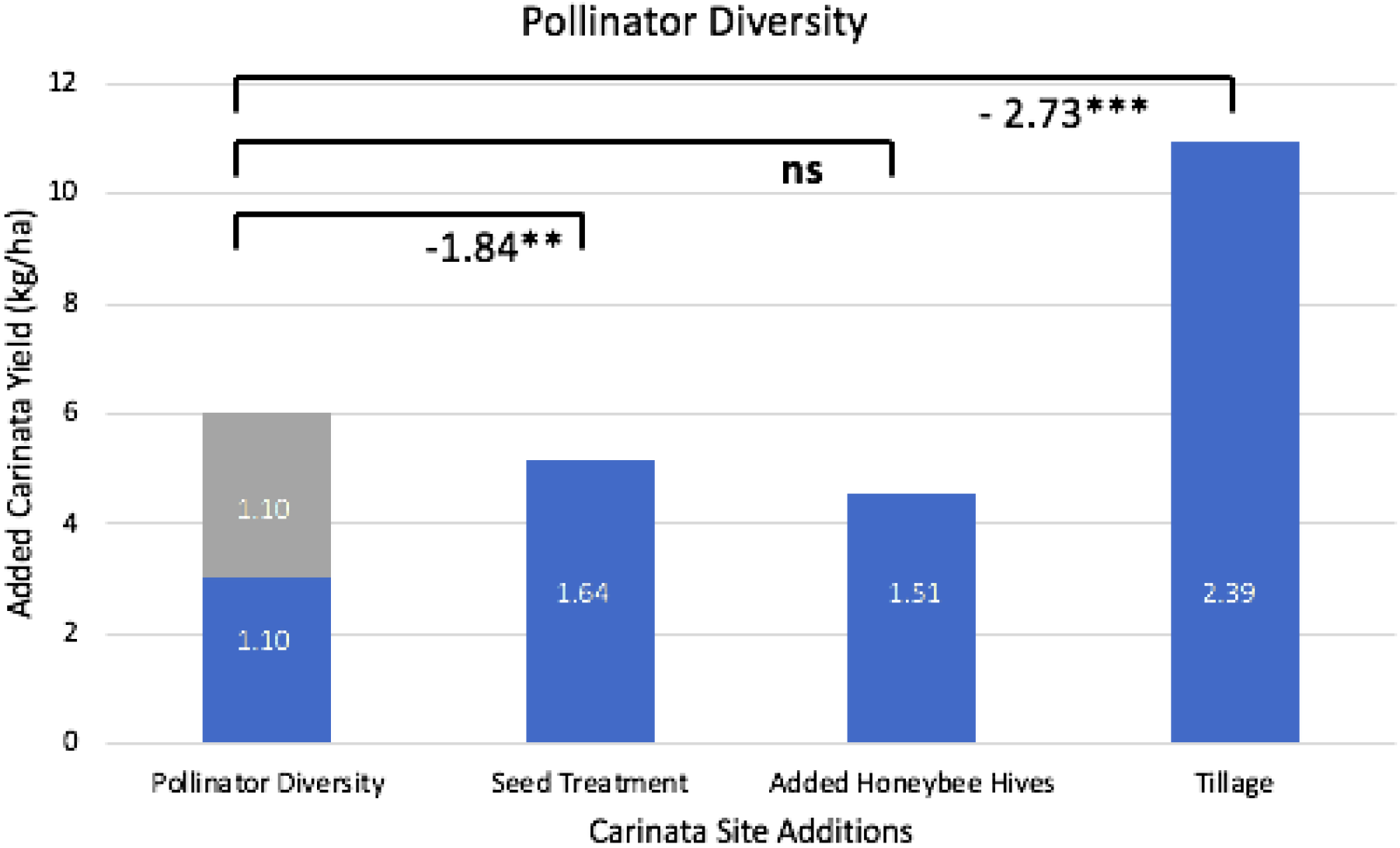
Best model estimates of carinata yield added by each farming practice and Shannon pollinator diversity shown as main effects. The estimates are based on the fitted values from the model including interactions of farming practices and insect diversity. The reduction or increase of yield depending on interactions with common farming practices is plotted in Fig 3. The blue Pollinator Diversity bar represents an increase from 1-2 on the Shannon index, while the gray bar represents a jump from 2-3. Numbers inside the bars represent effect sizes before back-transformation. Brackets indicate the negative interactions between insect diversity and the common farming practices. *P* < 0.05; * *P* < 0.05; ** *P* < 0.01; *** *P* < 0.001.

### Model 3) FARMING PRACTICES ON INSECT AND POLLINATOR DIVERSITY

Using insect and pollinator diversity as response variables and farming practices as predictor variables, we did not find significant evidence that any combination of farming practices within our one-acre sites enhances or decreases insect or pollinator diversity.

### Model 4) LANDSCAPE HETEROGENEITY AND INSECT DIVERSITY

#### Best model for insect diversity including all farming practices and landscape heterogeneity

The minimal adequate model includes two significant two-way interactions: There is a significant positive interaction between landscape heterogeneity at the 500 m scale and seed treatment with an effect of 0.15±0.07 (increasing insect diversity by 1.14 H) (LRT: χ^2^_1DF_ = 5.1146, p = 0.023) (Fig. 6, Appendix S3: Table S3). There is also a significant negative interaction between landscape heterogeneity at the 1000 m scale and tillage with an effect of - 0.18±0.08 (increasing diversity by 0.99 H) (LRT: χ^2^_1DF_ = 5.5933, p = 0.018) (Fig. 5, Appendix S3: Table S3).

**FIGURE 5:**
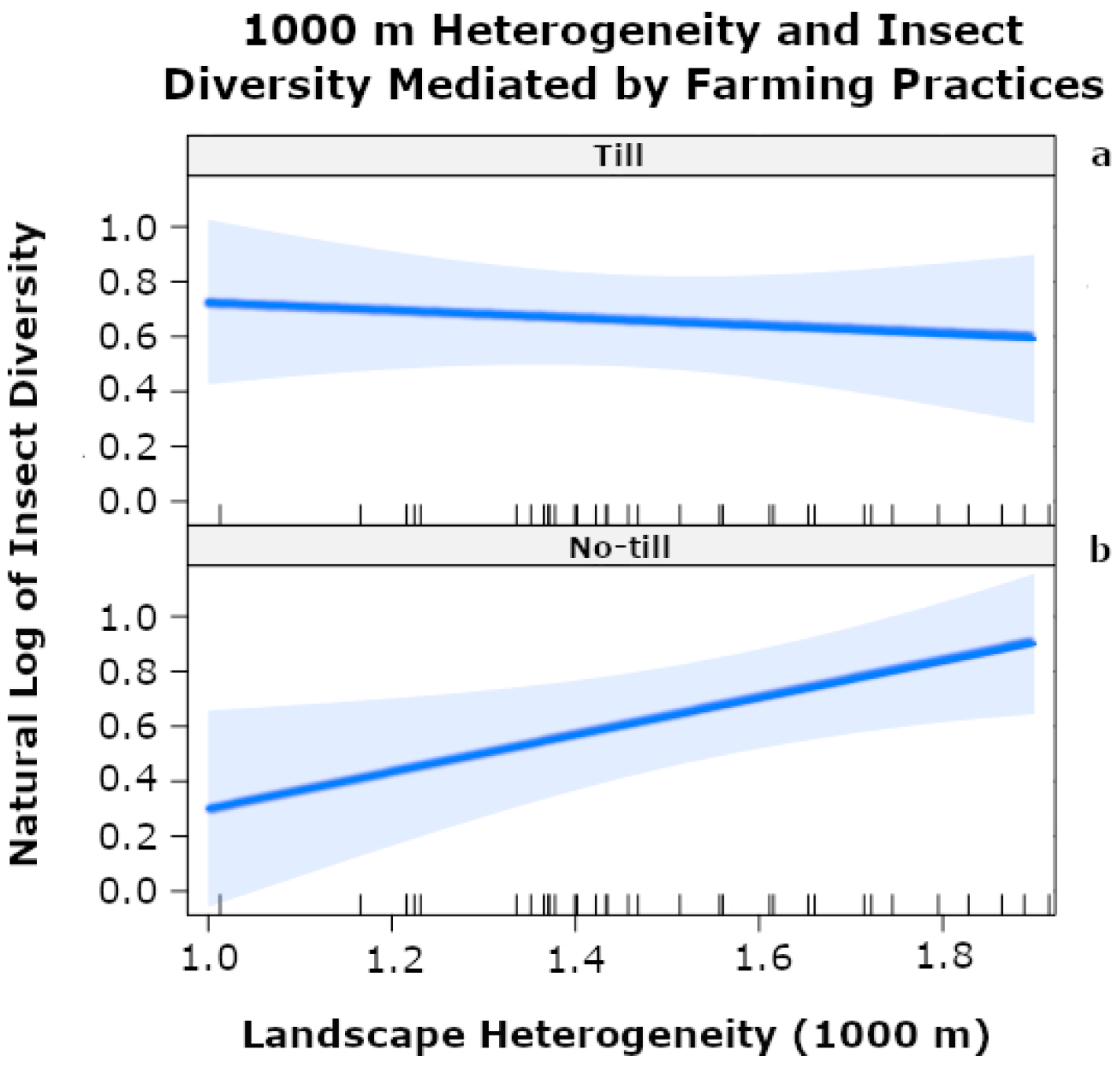
Linear mixed effect analysis of interactions between tillage and landscape heterogeneity at a 1000 m radius on insect diversity sampled within sites of *Brassica carinata* in eastern South Dakota, 2017-2018. Ticks on the x-axis represent measured individual site heterogeneity.

**FIGURE 6:**
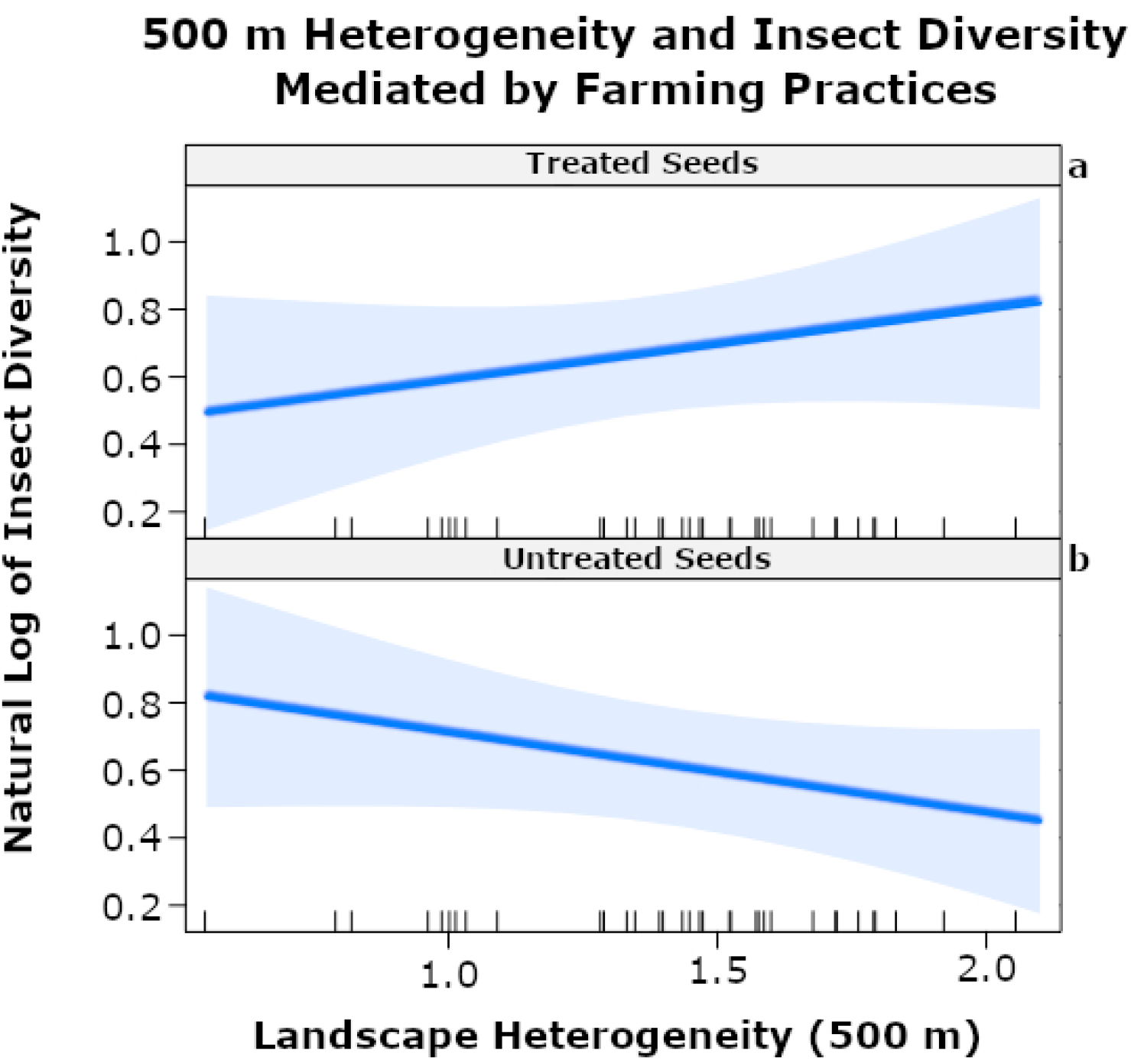
Linear mixed effect analysis of interactions between seed treatment and landscape heterogeneity at a 500 m radius on insect diversity sampled within sites of *Brassica carinata* in eastern South Dakota, 2017-2018. Ticks on the x-axis represent measured individual site heterogeneity.

#### Main effects of farming practices and landscape heterogeneity on insect diversity

There is a significant positive relationship between insect diversity and landscape heterogeneity at the 1000 m scale with an effect of 0.14±0.05 increasing insect diversity by 1.16 H (p = 0.019) (Fig. 5, Appendix S3: Table S3).

### Model 5) LANDSCAPE HETEROGENEITY ON POLLINATOR DIVERSITY

#### Best model for pollinator diversity including all farming practices and landscape heterogeneity

The minimal adequate model includes a significant two-way interaction. There is a significant negative interaction between landscape heterogeneity at the 3000 m scale and tillage on pollinator diversity with an effect of −0.39±0.1(0.94 H) (LRT: χ^2^_1DF_ = 12.47, p < 0.001) (Fig. 7, Appendix S3: Table S4).

**FIGURE 7:**
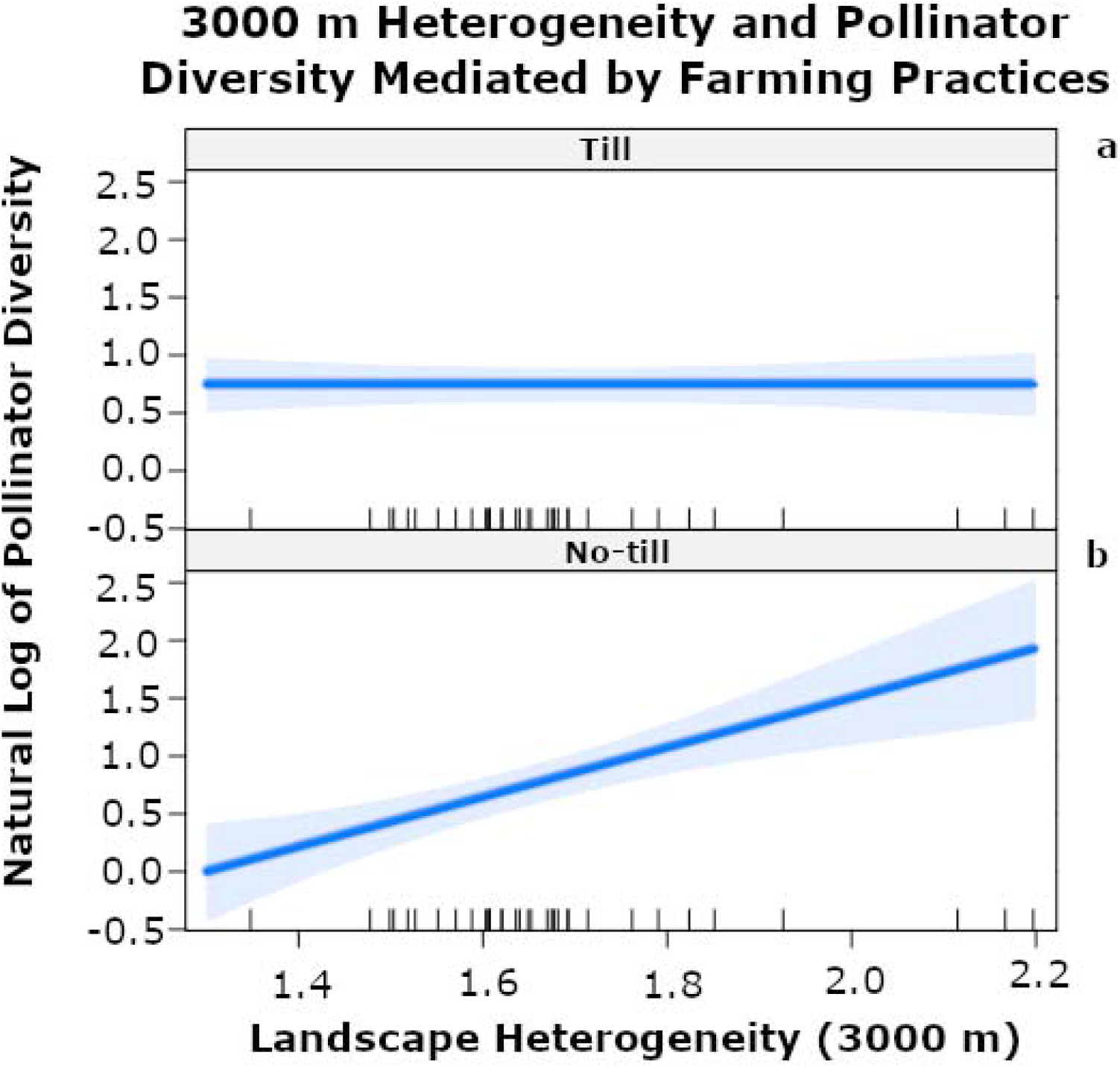
Linear mixed effect analysis of interactions between tillage and landscape heterogeneity at a 3000 m radius on pollinator diversity sampled within sites of *Brassica carinata* in eastern South Dakota, 2017-2018. Ticks on the x-axis represent measured individual site heterogeneity.

#### Main effects of farming practices and landscape heterogeneity on pollinator diversity

There is a significant positive relationship between landscape heterogeneity and pollinator diversity at the 3000 m scale with an effect of 0.39±0.09 (p < 0.001) (1.49 H) (Fig. 7, Appendix S3: Table S4).

### Model 6) LANDSCAPE HETEROGENEITY ON YIELD

#### Best model for yield including landscape heterogeneity and all farming practices

The minimal adequate model includes a significant two-way interaction. There is a significant negative interaction between landscape heterogeneity at the 3000 m scale and tillage with an effect of −1.85±0.5 (adding 2.44 kg/ha for every unit increase in heterogeneity) (LRT: χ^2^_1DF_ = 12.505, p < 0.001) (Appendix S3: Table S5).

#### Main effects of landscape heterogeneity and farming practices on yield

There is a significant positive relationship between landscape heterogeneity at the 3000 m scale and yield of carinata with an effect of 2.13±0.46 (p < 0.001) (adding 8.47±1.59 kg/ha for every unit increase in heterogeneity) (Appendix S3: Table S5).

## Discussion

Few studies have evaluated the relationship between overall insect/pollinator diversity and yield in the context of farming practices (Letourneau and Bothwell 2008, Lundgren and Fausti 2015). The approaches used here allow us to address the importance of three common farming practices and insect/pollinator diversity to yield, landscape heterogeneity to insect/pollinator diversity, and lastly, landscape heterogeneity to yield. Overall, we found that increased insect/pollinator diversity as well as the common farming practices (added honey bee hives, tillage, neonicotinoid seed treatment) increase carinata yield. However, the common farming practices studied interfere with pollination services and natural pest control services provided by wild insects, perhaps through competition with native insects in the case of added honey bee hives or by killing beneficial insects in the case of seed treatment and tillage. Additionally, we found that insect/pollinator diversity within our carinata sites is dependent on large-scale landscape heterogeneity and not on farming practices within our sites. Finally, we demonstrate that the largest scale of landscape heterogeneity (3000 m) is positively related to carinata yield. Below we address in turn the ecosystem services provided by insect and pollinator diversity to carinata yield, and the landscape determinants affecting insect and pollinator diversity.

### Insect/pollinator diversity effects on carinata yield

A single unit increase of insect diversity had a stronger positive effect on yield than did treating crops with neonicotinoids. Our results are consistent with previous findings that biodiversity and ecosystem services are positively related (Chapin et al. 2000, Hooper et al. 2005) and higher levels of pollinator diversity are associated with increased yield (Greenleaf and Kremen 2006, Atmowidi et al. 2007, Hoehn et al. 2008, Mallinger and Gratton 2015, Dainese et al. 2019).

Increases in insect diversity and the addition of honey bees had equal effects on the yield Insect diversity has a stronger effect size (2.28) than pollinator diversity (1.10) on yield, indicating that insect diversity contributes more to a higher yield than does pollinator diversity alone. Insect diversity includes all pollinators collected, but it also accounts for non-pollinating insects including both pests, predators, and parasitoids. Insect diversity could favor suppression of pest populations and enhance the activity of predators and parasitoids in agroecosystems (Landis et al. 2000), especially if the measured biodiversity exists within a complex landscape (Bianchi et al. 2006). Therefore, many non-pollinating insects also contribute ecosystem services by suppressing pest populations (natural pest control). Our insect diversity measurement more strongly contributes to carinata yield our pollinator diversity measurement because it includes pollinators, predators, and parasitoids. We observed several commonly known predators of pests such as parasitic wasps (Hymenoptera: Ichneumonidae and Braconidae), lacewings (Neuroptera), lady beetles (Coleoptera: Coccinellidae), and hemipteran predators in the family Reduviidae, all of which could contribute to the decrease in pest abundance.

### Insect/pollinator effects on yield are modified by common farming practices

Some crops do not show yield increases with neonicotinoid treatment (Seagraves and Lundgren 2012). However, neonicotinoids, tillage (Malhi and Lemke 2007), and added honey bee hives (Sabbahi et al. 2005), have all been observed to increase yields of canola, and these observations are congruent with our results with the relative carinata. The addition of any single farming practice to the relationship between yield and insect/pollinator diversity is negative, indicating that farming practices interfere with the positive effects on yield by insect/pollinator diversity. Tilling almost completely negated the positive effect of insect or pollinator diversity on yield. These findings are consistent with previous studies demonstrating that eusocial bees are more sensitive to tilling regardless of nest location (Williams et al. 2010, Kratschmer et al. 2018). A single fertile female is responsible for eusocial bee reproduction, which could lead to greater difficulty in repopulation after disturbance compared to solitary species in which most females are reproductive (Kratschmer et al. 2018). There is also a negative effect of seed treatment with insect/pollinator diversity in relation to yield, supporting previous studies that even sublethal doses of neonicotinoids negatively alter pollinator behavior (Henry et al. 2012, Rundlöf et al. 2015). This suggests that seed treatment decouples the relationship between insect/pollinator diversity and yield. There is also negative interaction between added honey bee hives and insect diversity with yield. There is no interaction between pollinator diversity and added honey bee hives, however. There is a possibility that non-pollinating insects could consume floral resources, contributing to the negative interaction between insect diversity and honey bee hives. Bees will avoid flowers containing predators and also flowers in which a previous predation attempt occurred (Dukas 2001). Furthermore, honey bees are competitive against other wild pollinators but are less efficient (Lindström et al. 2016) potentially resulting in lower yield. This could explain the negative interaction that honey bees have indirectly on yield by competing with wild pollinators and avoiding some flowers that are occupied by non-pollinating insects. However, more research is needed to determine the mechanisms underlying the relationship between honey bees and non-pollinating insects.

The lack of an interaction between pollinator diversity and added honey bee hives suggests that the effects of wild pollinators and honey bees are additive. The foraging behavior and size of the honey bees are unique among observed pollinators in the field, possibly filling a niche in carinata pollination requirements. Bee species often have differing forage heights, time of day, and behavior on the flower (Hoehn et al. 2008), suggesting that diverse preferences are the mechanism by which species diversity operates. In particular, pollinator preferences depends on the spatiotemporal availability of floral resources (Nottebrock et al. 2017). Pollinators more frequently visit plants with high amounts of nectar (Schmid et al. 2016), which especially counts for honey bees that are more abundant on sites with mass flowering crops (Holzschuh et al. 2013). Honey bees could therefore supplement yield provided by wild bees (Garibaldi et al. 2013) maximizing the yield of a site. This is especially important when considering recent declines in wild bee diversity and abundance (Burkle et al. 2013). Current research on competitive interactions between managed honey bees and native bees is mixed, with about half of studies finding negative interactions between managed and native bees (Mallinger et al. 2017). In our study, we are able to show that honey bees are more important on sites that have been tilled but are problematic on sites with high insect diversity and no soil tillage and pesticide treatment.

Four three-way interactions were observed within our models. Two occurred in our model relating to pollinators and yield (seed treatment × tillage × pollinator diversity and added honey bee hives × tillage × pollinator diversity) while two occurred in our insect and yield model (seed treatment × added honey bee hives × insect diversity and seed treatment × tillage × insect diversity), all were positive. All interactions contained to a diversity metric and two common farming practices. It is possible that yield is compensated by the addition of a second farming practice to offset the negative interaction between a single farming practice and a diversity metric on yield. One farming practice might interact negatively with insect/pollinator diversity and require an additional farming practice to compensate for those losses in yield. Mechanisms behind these interactions are not apparent, but deserve more attention. There is no interaction between all three farming practices with insect/pollinator diversity, suggesting that these insect/pollinator and yield relationships reach a threshold in which they are not altered any further.

Our findings demonstrate that benefits provided by a diverse insect/pollinator community can be decoupled by human modification of the landscape. Stakeholders should be cautious before intensive farming practices are implemented. We conclude that the farming practices manipulated in our study negatively alter the ecosystem services provided by insects; thus, producing a cap on how much yield can be attained on a specific field when different farming practices are performed.

### Landscape effects on insect/pollinator diversity

There was no effect of small-scale landscape heterogeneity (500 m) on yield or insect diversity, demonstrating that large-scale land use (>500 m) is important for insect/pollinator diversity and that biodiversity loss associated with land use change is not likely an issue that can be addressed by a single landowner. Honey bees forage up to 6 km (Beekman and Ratnieks 2000) and bumble bees can fly up to 20 km (Morris 1993). As landscape heterogeneity increases, the functional land uses available to insects will also increase, such as forage for resources, nesting, and mating grounds (Fahrig et al. 2011). Landscape heterogeneity at the 1000 m scale was significant to total insect diversity because many non-pollinating insects are not as strong of fliers and cannot travel the same distances as pollinators. Yield was significantly positively associated with landscape heterogeneity at the 3000 m scale. Because heterogeneity at the same time is positively related to pollinator diversity and yield at 3000m scale, we conclude that pollinator diversity is enhanced by landscape heterogeneity. A more heterogenous landscape may reflect farming practices that positively influence insect diversity and edge area between land uses, both of which contribute to yield.

### Landscape effects on insect/pollinator diversity are modified by common farming practices

The significant positive interaction between small scale landscape heterogeneity and seed treatment on insect diversity could be related to the behavioral alterations of neonicotinoids on insect behavior (Tomé et al. 2012, Williamson et al. 2014) including a developed preference for neonicotinoid laced resources (Arce et al. 2018). Thus, a treated site could capture more species from the diverse local landscape. Alternatively, insects present in neonicotinoid treated sites could be paralyzed and less capable of long distance travel. Thus, we capture a greater diversity of insects during any observational period. The negative interaction between 1000 m landscape heterogeneity and 3000 m landscape heterogeneity with tilling in relation to insect and pollinator diversity may indicate that tillage practices destroy the habitat suitability for insects and pollinators (Nicholls and Altieri 2013). Tilling at our carinata plots almost completely negated the positive effects of a heterogeneous landscape on insect/pollinator diversity. Increased heterogeneity provides a greater diversity of functional habitats for insects and pollinators, (Steffan-Dewenter 2002, Tscharntke et al. 2005) many of which are ground nesting and would be destroyed by tillage (Kratschmer et al. 2018).

## Conclusions

Carinata yield at our plots is increased by 1) common farming practices (neonicotinoid treatment, tillage, and added honey bee hives), 2) increasing diversity of pollinating insects, 3) increasing diversity of the entire insect community and 4) increasing landscape heterogeneity. There is, however, tension between many of the farming practices and insect/pollinator diversity. Many farming practices might ultimately increase yield, but they are not additive in that they decrease the insect/pollinator contribution to yield. Our findings suggest that increased landscape heterogeneity and insect/pollinator diversity increases yield, but these relationships are decoupled by common farming practices such as tilling, seed treatment, and added honey bee hives. Additionally, mass flowering crops could increase the abundance of wild bee species (Holzschuh et al. 2013) but the studied bee was a solitary species and might not reflect the behavior of eusocial species. These crops could be a way to sustain native pollinators without inflicting severe economic harm on producers.

Human land use does not necessarily entail habitat destruction, and proper agricultural management can enhance biodiversity, in turn increasing ecosystem function and services (Tscharntke et al. 2005). Management tactics such as diversification of the landscape, reduction in tilling, and reduction of pesticide use could all have positive impacts on the ecosystem services provided by pollinators. Increased landscape heterogeneity increases biodiversity and will therefore, act as biological insurance for ecosystem services (Loreau et al. 2003). This study could have policy implications relating to the use of pesticides, tilling, and the diversification of the landscape. Policies that discourage the use of tilling and pesticides could be paired with incentives to diversify the landscape, maximizing pollinator health in an agricultural landscape. The warming of the global climate is predicted to increase arable land in North America by 40% (Fischer et al. 2005). By providing insects with a diverse and connected landscape we can invest in the future of agriculture in the US. Climate change in northern areas, such as the Upper Midwest, could allow for the introduction of new crop species (Olesen and Bindi 2002). A diverse and healthy array of insects could be maintained in a heterogenous landscape, maximizing the benefits this climate change could bring. Our results demonstrate that landscape heterogeneity is an important factor in the enhancement of pollination services, that not only increase yield, but could allow us to accommodate crops suitable to the future climate of this region.

## Acknowledgements

We thank I. Vilella-Arnizaut, N. Petersen, J. Gelderman, and J. Smithers for their assistance in the field and lab. M. Latvis provided conceptual insight. This research was funded by the North Central Sun Grant Initiative (USDA/DOE) SA1500640.

